# The prelimbic prefrontal cortex mediates the development of lasting social phobia as a consequence of social threat conditioning

**DOI:** 10.1101/2024.06.04.597446

**Authors:** Kelly Lozano-Ortiz, Ada C. Felix-Ortiz, Jaelyn M. Terrell, Angelica R. Ramos, Jose Rodriguez-Romaguera, Anthony Burgos-Robles

## Abstract

Social phobia is highly detrimental for social behavior, mental health, and productivity. Despite much previous research, the behavioral and neurobiological mechanisms associated with the development of social phobia remain elusive. To investigate these issues, the present study implemented a mouse model of social threat conditioning in which mice received electric shock punishment upon interactions with unfamiliar conspecifics. This resulted in immediate reductions in social behavior and robust increases in defensive mechanisms such as avoidance, freezing, darting, and ambivalent stretched posture. Furthermore, social deficits lasted for prolonged periods and were independent of contextual settings, sex variables, or particular identity of the social stimuli. Shedding new light into the neurobiological factors contributing to this phenomenon, we found that optogenetic silencing of the prelimbic (PL), but not the infralimbic (IL), subregion of the medial prefrontal cortex (mPFC) during training led to subsequent forgetting and development of lasting social phobia. Similarly, pharmacological inhibition of NMDARs in PL also impaired the development of social phobia. These findings are consistent with the notion that social-related trauma is a prominent risk factor for the development of social phobia, and that this phenomenon engages learning-related mechanisms within the prelimbic prefrontal cortex to promote prolonged representations of social threat.

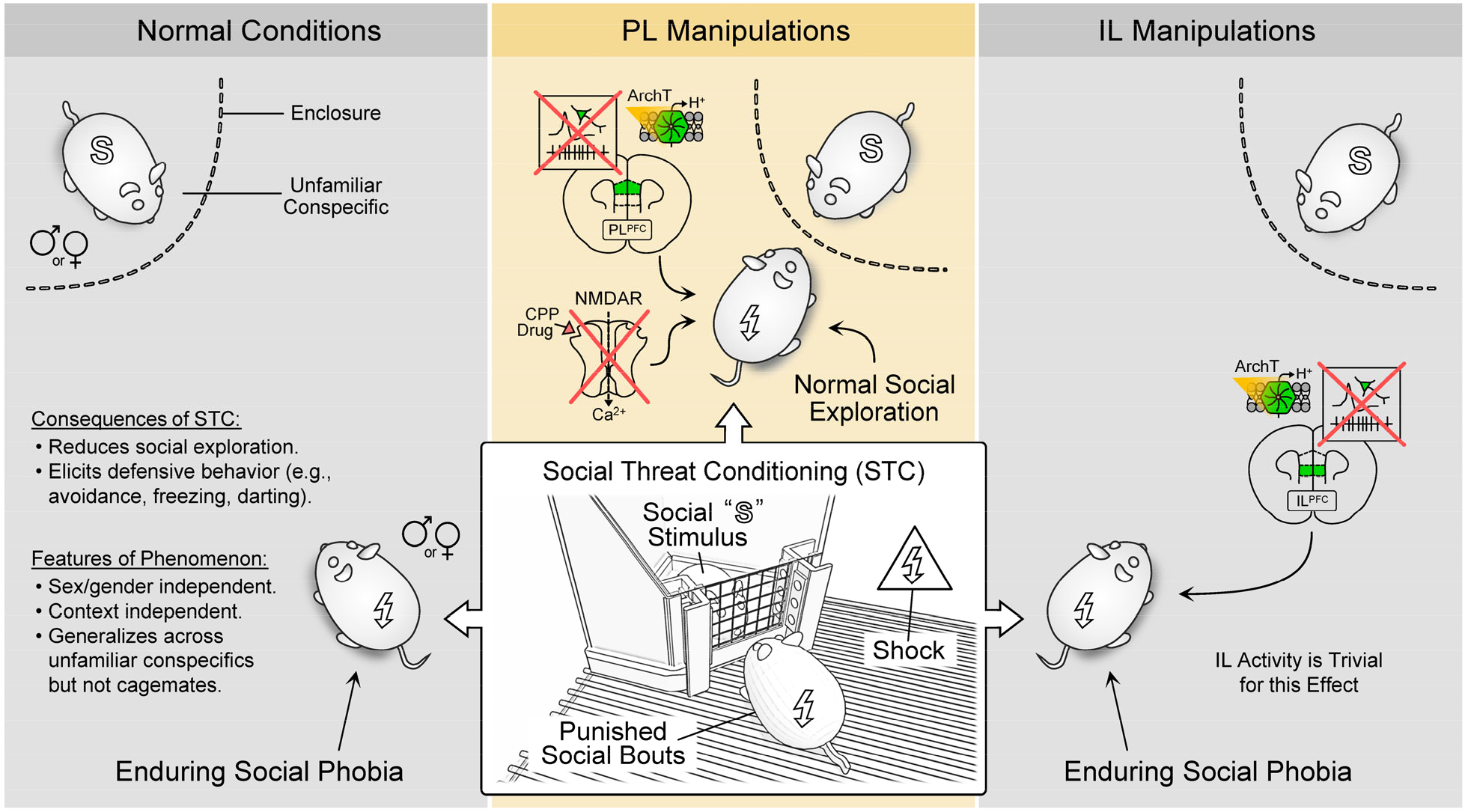

## INTRODUCTION

Social phobia is an increasingly prevalent anxiety-related disorder that affects up to 12% of individuals [1–4]. Such high incidence keeps prompting new research to better understand the mechanisms, etiology, and most critical factors contributing to this condition [5–8]. In addition to research in human patients [9,10], scientists have developed multiple animal models that are very useful to study social phobia [11–14]. One such model is social threat conditioning (STC), which focuses on the negative impacts of socially-derived traumatic experiences on social behavior [15,16]. Yet, this paradigm remains sparsely studied, and thus, many critical gaps still exist in knowledge regarding this phenomenon.

STC is based on principles from operant threat conditioning. Briefly, in this paradigm the animal subjects receive electric shock punishment upon interactions with unfamiliar conspecifics. This produces robust impairments in social behavior that persist for prolonged periods, even if the animal subjects are tested against new unfamiliar conspecifics [15,16]. While these observations are consistent with induction of phobia-like states, the following questions still remain elusive. Are impacts on social behavior the only indicators of social phobia during STC, or are there other behavioral indicators of social threat learning during this paradigm? Could stimulus specificity be achieved during this paradigm, or does social stimulus generalization always prevail after STC? Is STC influenced by sex differences? Finally, are there any significant sex differences when this phenomenon occurs with matched or mismatched sexes between the test subjects and the social predictors of threat?

Regarding neurobiological substrates, social phobia has been associated with hyperfunction of the medial prefrontal cortex (mPFC). In human patients, dorsal regions of the mPFC exhibit increased cortical thickness, elevated baseline activity, and augmented responsiveness during the presentation of social stimuli [17–20]. Studies in rodents have also implicated mPFC areas during behavioral manifestations of social phobia. For instance, in a study that implemented a social defeat paradigm in mice, eventual reductions in social behavior were associated with increased cfos expression in the mPFC [21]. Consistently, in a study that used the STC paradigm, subsequent reductions in social behavior were attributed to increased neuronal activity in mPFC [22]. While these observations implicate the mPFC in the expression of social phobia, a direct link between mPFC activity during social-related trauma and the eventual development of social phobia has not been established yet.

Using an improved version of the paradigm, the present study elucidated multiple new features of STC, including impacts on anxiety-like avoidance and other defensive behaviors, and addressed issues on stimulus specificity versus generalization, as well as on potential sex differences. Optogenetic silencing and drug microinfusion approaches were then implemented to determine whether neural activity and plasticity-related molecular events are crucial mechanisms by which distinct subregions of the mPFC (PL and IL) promote the encoding of prolonged representations of social threat. Impacts on various behaviors were evaluated throughout the study, including impacts on social behavior, avoidance, freezing, darting, and stretched posture, all of which represent arousal states and defensive mechanisms during threat [23–25]. Overall, this broader approach provided novel insights and previously unreported perspectives on the mechanisms contributing to social threat conditioning.

## MATERIALS AND METHODS

### Animal Subjects

All procedures were approved by the IACUC in compliance with U.S. PHS Policy and The Guide. A total of 194 wild-type C57BL/6J mice (including males and females; Jackson Laboratory) were used in this study (138 served as test subjects, 48 served as social stimuli, and 8 served as naïve controls). Mice were ∼10 weeks old at arrival and were group housed (4 per cage) for the entire length of experiments. The animal vivarium had controlled temperature, pressure, and a 12-hr light/dark regular cycle with lights on at 7:00 A.M. Food and water were available ad libitum.

### Social Threat Conditioning

STC, also known as social fear conditioning, was performed using principles from previous studies [11,15,22]. Yet, significant improvements were made for broader behavioral analysis and full automation of the task. The task was conducted in standard conditioning chambers (32×25×22 cm) that were equipped with grid floors and current scramblers for footshocks (Med Associates). Two smaller customized enclosures were included in each chamber for presentations of social stimuli (i.e., conspecifics of the same age and weight as the test subjects). These enclosures were developed using conventional 3D printing procedures, and were fitted with acrylic panels and stainless-steel mesh screens through which mice could interact with each other. Social interactions were detected using photodetectors that were mounted on both sides of the mesh screens (*more details in the supplementary materials*). During the initial 5-min acclimation period, the test subjects were allowed to freely explore the chambers, enclosures, and social stimuli without any consequence. During the subsequent 20-min conditioning period, social bouts that lasted >1 s were paired with footshock (0.40 mA, 0.5 s) to the test subject.

### Other Behavioral Assays

Sociability tests were conducted before and after STC to evaluate impacts on multiple behaviors. These tests were conducted in rectangular acrylic boxes (50×30×22 cm), each including two inverted wire-mesh cups (Ø9.5 cm, 2-mm openings) in the central zone. In most experiments, one cup contained a social stimulus, while the other cup remained empty, otherwise indicated. The social stimuli were rotated on a daily basis, so that the test subjects were always tested against new unfamiliar conspecifics, otherwise indicated. At the beginning of each session, the test subjects were placed in the middle of the arena and allowed to freely explore the apparatus, cups, and social stimuli for a total of 15 min.

Open field tests were also conducted to evaluate possible effects in general anxiety and locomotion [26,27]. These were conducted in squared acrylic boxes (40×40×40 cm). The arena was virtually divided into a peripheral zone and a central zone (24×24 cm). At the beginning of the open field tests, mice were placed in the center zone and allowed to freely explore the apparatus for a total of 9 min. During open field tests that involved optogenetic manipulations, the assay was divided into 3-min epochs with alternating laser status (Off-On-Off) to evaluate potential transitions in behavioral effects.

### Behavioral Analysis

ANY-maze software was used to control the hardware, record videos at 10 fps, and quantify most behaviors in an automated manner. The time that the test subjects spent in different zones was quantified as proxies for (a) social exploration, (b) non-social exploration, and (c) anxiety-related avoidance. Additional quantifications were extracted from ANY-maze, including: (a) fear-related freezing, (b) number of social bouts, (c) number of shocks, (d) distance to the stimulus, (e) number of entries, (f) total distance traveled, and (g) locomotion speed. Hand-scoring methods were also implemented by experimenters who remained blind to treatment to evaluate additional behaviors, such as: (a) forward darting, (b) reversed darting, (c) stimulus-oriented darting, and (d) stretching postures. More details on behavioral quantifications, data analyses, and statistical tests are provided in the supplementary materials.

### Stereotaxic Surgeries

Standard surgical procedures were used for viral infusions and chronic implantation of optical fibers or cannulas [28]. Details on surgical procedures, equipment, anesthesia, analgesia, stereotaxic coordinates, and post-operative care are provided in the supplementary methods.

### Optogenetic Silencing

Standard procedures were used for optogenetic silencing of neuronal activity. While controls only expressed enhanced yellow fluorescent protein (eYFP), the neural silencing groups expressed archaerhodopsin (ArchT), which was triggered using red-shifted DPSS lasers (589-nm; OptoEngine). Laser light was delivered through chronically implanted optical fibers, which were connected to patch cords (1×2, Ø200-µm core, FCM-2xMF1.25; Doric Lenses) and fiberoptic rotary joints (FRJ-1×1; Doric Lenses) for free movement. Light was delivered in a constant fashion (∼5 mW, ∼18 mW/mm^2^) as mice underwent the STC task. More details are provided in the supplementary methods.

### Drug Microinfusions

Standard procedures were used for drug microinfusions via chronically implanted bilateral cannulas [29]. Mice received either artificial cerebrospinal fluid (ACSF) or a long-acting competitive antagonist of NMDARs (CPP, 50 ng in 300 nL per side). More details can be found in the supplementary methods.

### Histology

Standard procedures were used to euthanize the test subjects, collect brains, and take images of the mPFC. Evaluation of viral expression, optical fiber placements, cannula positions, and drug microinfusion sites was then performed. Details for these procedures are provided in the supplementary methods.

## RESULTS

### Social threat conditioning produced robust behavioral traits of social phobia

After developing new hardware to perform STC in a fully automated manner (Supp Fig 1A-B), our first goal was to evaluate novel behavioral features during the STC task. On the first day of the experiment, two groups of male mice underwent a pre-conditioning sociability test in the presence of unfamiliar conspecifics to establish the baseline for multiple behaviors (Fig 1A). Quantifications of time spent in distinct zones of the apparatus allowed us to assess multiple behaviors in a systematic manner using software (Supp Fig 1C-D). While social exploration was the most prominent behavior during the baseline test, no significant differences were detected between the two groups tested (Fig 1B-C). Group differences later on were then attributed to distinct experiences during the STC task.

**Fig 1.**
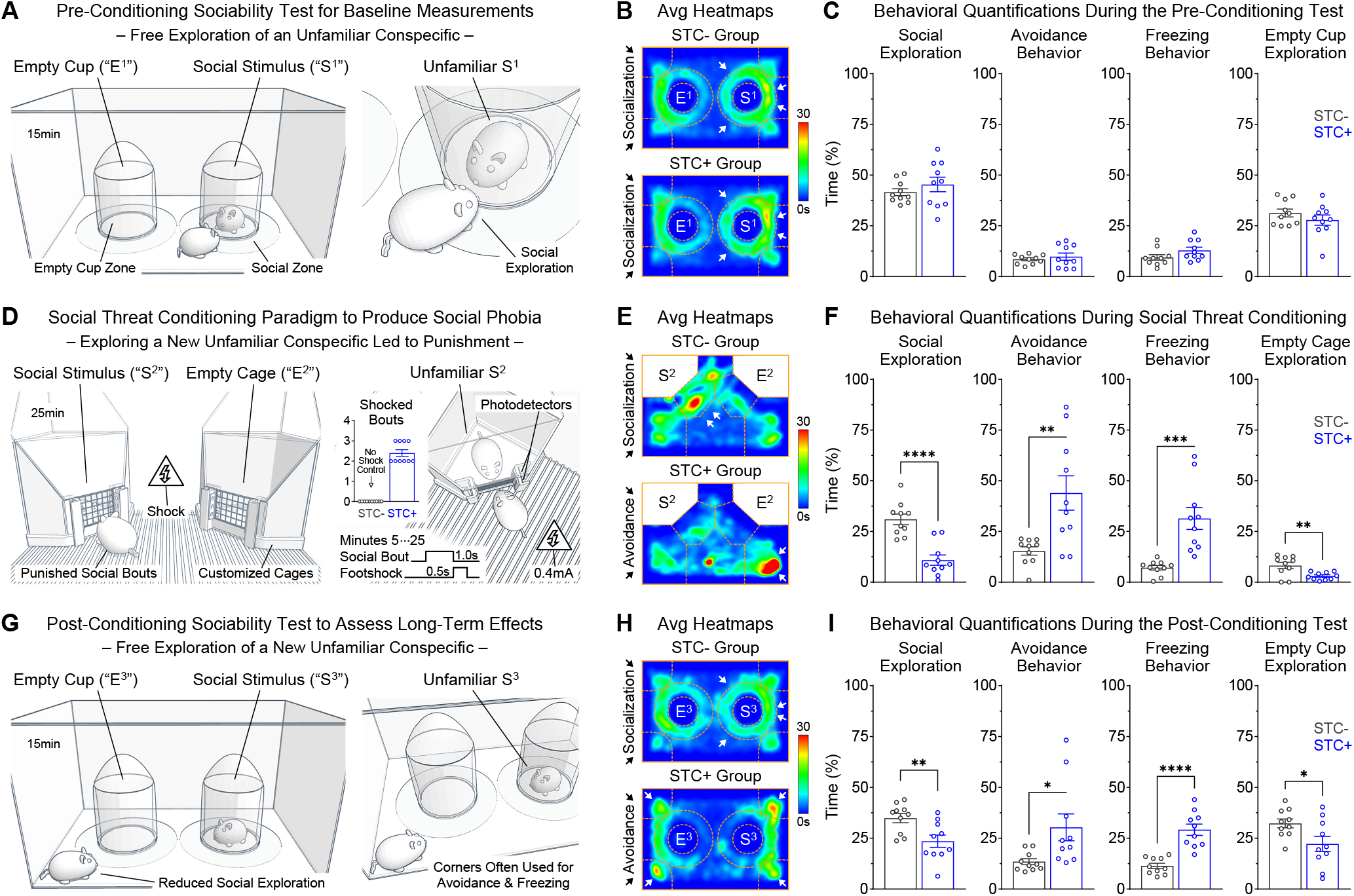
Social threat conditioning produced a behavioral profile of social phobia. ***A-C***, Pre-conditioning sociability test to establish behavioral baselines. Social exploration was quantified as time spent in proximity to the cup containing an unfamiliar conspecific. Avoidance was quantified as time spent in the corners of the box. Freezing was quantified as time of complete immobilization anywhere in the box. Social exploration was the most prominent behavior (Social Cup, *t*_(18)_ = 0.97, *P* = 0.34; Avoidance, *t*_(18)_ = 0.69, *P* = 0.50; Freezing, *t*_(18)_ = 1.78, *P* = 0.093; Empty Cup, *t*_(18)_ = 1.09, *P* = 0.29). ***D-F***, Social threat conditioning twenty-four hours after the baseline test. During the first 5 min of acclimation, both groups were allowed to freely explore a new unfamiliar conspecific. During the subsequent 20 min, while the control group (STC-, N = 10 males) continued exploring the unfamiliar conspecific without any consequence, the experimental group (STC+, N = 10 males) received electric shocks during social bouts. This produced decreases in social exploration and increases in avoidance and freezing (Social Enclosure, *t*_(18)_ = 5.51, *P* < 0.0001; Avoidance, *t*_(18)_ = 3.27, *P* = 0.0042; Freezing, *t*_(18)_ = 4.38, *P* = 0.0004; Empty Enclosure, *t*_(18)_ = 3.09, *P* = 0.0064). ***G-I***, Post-conditioning sociability test twenty-four hours after conditioning. Regardless of the new unfamiliar conspecific and a context that differed from the conditioning context, the STC+ group exhibited a behavioral profile of social phobia (Social Cup, *t*_(18)_ = 2.99, *P* = 0.0079; Avoidance, *t*_(18)_ = 2.48, *P* = 0.023; Freezing, *t*_(18)_ = 5.83, *P* < 0.0001; Empty Cup, *t*_(18)_ = 2.33, *P* = 0.032). [In all figures, data is illustrated as mean ± sem with superimposed values for individual subjects. \**P* < 0.05, \*\**P* < 0.01, \*\*\**P* < 0.001, \*\*\*\**P* < 0.0001]

On the second day of the experiment, the two groups were introduced to the conditioning chambers, each containing two novel enclosures, one of which contained a new unfamiliar conspecific. After a brief acclimation period, the control group (STC-, N = 10 males) continued exploring the unfamiliar conspecific without any consequence. In contrast, the experimental group (STC+, N = 10 male) received footshocks every time they interacted with the unfamiliar conspecific (Fig 1D). After a few punished social bouts, the STC+ group exhibited drastic shifts in behavior that were characterized by significant reductions in social exploration and significant elevations in avoidance and freezing, compared to the STC-controls (Fig 1E-F). Upon visual inspection of representative videos, it was evident that other defensive mechanisms also emerged during the STC task (Supp Fig 2). These included stretched postures which are associated with risk assessment and avoidance-approach ambivalence [30,31], and darting moves which represent escape and flight-like responses [32,33]. These findings are consistent with the effectiveness of the STC task to produce social phobia states.

**Fig 2.**
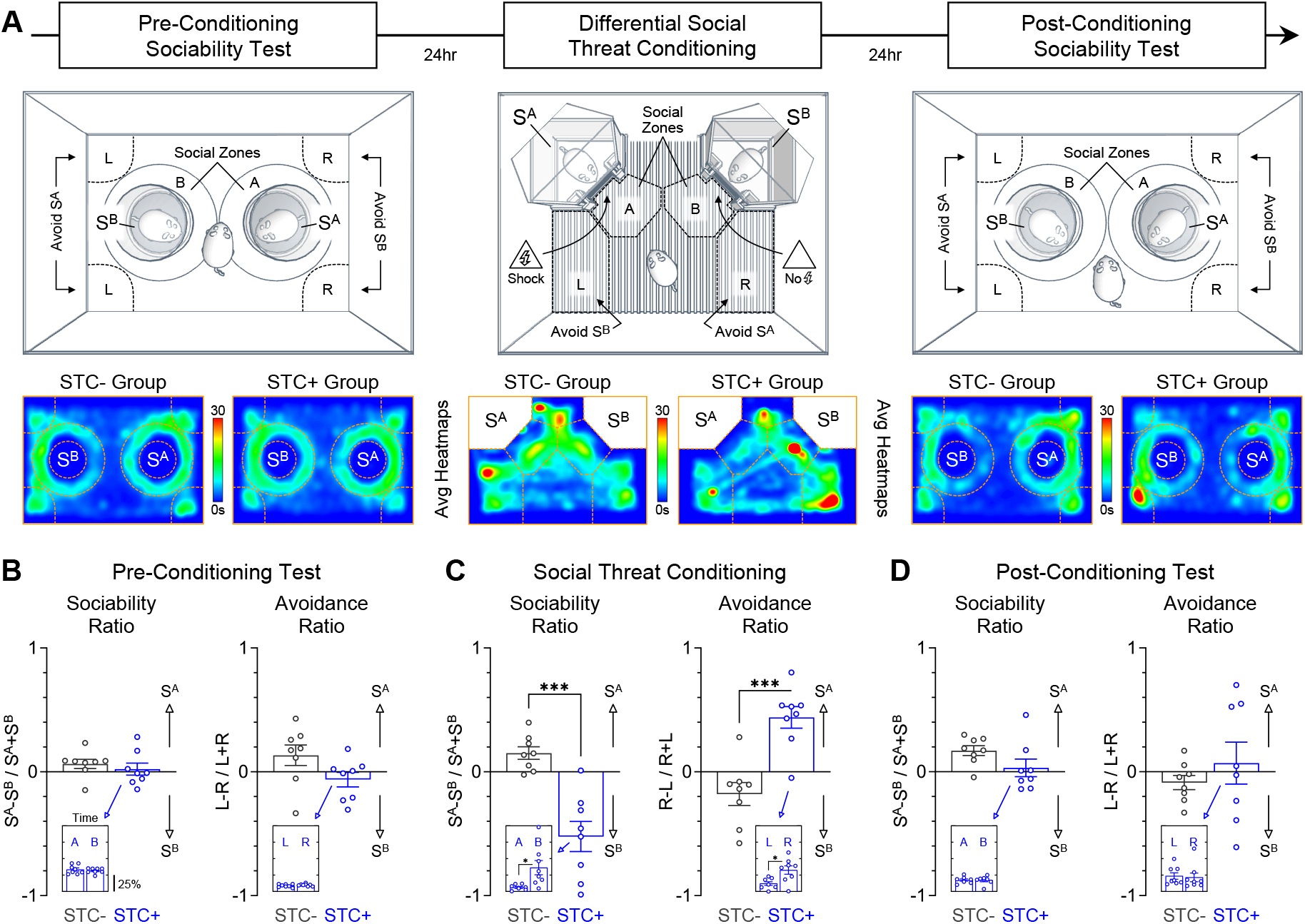
Differential social threat conditioning also resulted in subsequent generalization across stimuli. ***A***, Modification of the paradigm to include two social stimuli (S^A^ and S^B^) simultaneously, but with only one getting paired with shock punishment (STC-, N = 8 males; STC+, N = 8 males). ***B***, Prior to conditioning, both groups exhibited similar sociability and avoidance levels to S^A^ and S^B^, which resulted in behavioral ratios that tended to zero in both groups (Sociability Ratio, *t*_(14)_ = 0.69, *P* = 0.50; Avoidance Ratio, *t*_(14)_ = 1.92, *P* = 0.075; Sociability Inset, *t*_(14)_ = 0.66, *P* = 0.52; Avoidance Inset, *t*_(14)_ = 1.03, *P* = 0.32). ***C***, During differential conditioning, while the STC-group continued to exhibit behavioral ratios that tended to zero, the STC+ group exhibited greater sociability with S^B^ which did not predict shock, and greater avoidance of S^A^ which predicted shock (Sociability Ratio, *t*_(14)_ = 5.12, *P* = 0.0002; Avoidance Ratio, *t*_(14)_ = 4.86, *P* = 0.0003; Sociability Inset, *t*_(14)_ = 2.78, *P* = 0.015; Avoidance Inset, *t*_(14)_ = 2.94, *P* = 0.011). ***D***, Despite successful differential conditioning the previous day, during the long-term test the STC+ group exhibited similar levels of sociability and avoidance to S^A^ and S^B^, which resulted in behavioral ratios that once again tended towards zero (Sociability Ratio, *t*_(14)_ = 1.73, *P* = 0.11; Avoidance Ratio, *t*_(14)_ = 0.88, *P* = 0.39; Sociability Inset, *t*_(14)_ < 0.01, *P* > 0.99; Avoidance Inset, *t*_(14)_ = 0.25, *P* = 0.80). [Insets show data as %time. \**P* < 0.05, \*\*\**P* < 0.001]

On the third day of the experiment, the two groups went through a new sociability test to evaluate the long-term effects of STC (Fig 1G). This test was conducted against new unfamiliar conspecifics, in a contextual setting that significantly differed from the conditioning context. Despite these new conditions, the STC+ group exhibited significantly lower social exploration and significantly higher avoidance and freezing behavior, compared to the STC-controls (Fig 1H-I). These results are consistent with the notion that STC produces robust behavioral shifts that resemble social phobia [15,22].

### Differential social threat conditioning still resulted in generalized social phobia

Our next goal was to test whether stimulus specificity could be achieved, using a new version of the STC paradigm in which two social stimuli were simultaneously presented, but only one became the predictor of shocks (i.e., differential social threat conditioning; Fig 2A). During the pre-conditioning test, the STC- and STC+ groups (N = 8 males per group) exhibited similar sociability with the two social stimuli (Fig 2B, Sociability Inset), and low avoidance of both stimuli (Fig 2B, Avoidance Inset). This resulted in balanced behavioral ratios and no differences between the groups (Fig 2B, Ratios). During differential conditioning, since the S^A^ stimulus predicted shocks, the STC+ group exhibited significantly lower sociability with S^A^ than with S^B^ (Fig 2C, Sociability Inset) and greater avoidance of S^A^ than S^B^ (Fig 2C, Avoidance Inset). This resulted in unbalanced behavioral ratios and significant differences between the groups (Fig 2C, Ratios). During the post-conditioning test, despite effective stimulus differentiation the previous day, the STC+ group showed low sociability with both S^A^ and S^B^ (Fig 2D, Sociability Inset), and avoidance oriented towards both stimuli (Fig 2D, Avoidance Inset). This resulted in balanced behavioral ratios and no differences between the STC- and STC+ groups (Fig 2D, Ratios). These findings suggest that stimulus specificity, or the particular identity of the social predictor of threat, does not necessarily play an important role during the STC task to produce subsequent deficits in social behavior. Instead, the STC task seems to produce generalization of social aversiveness, which reinforces the notion that this paradigm is ideal to evaluate the mechanisms contributing to social phobia [11,15,16].

### Social deficits were persistent, independent of sex variables, and specific to unfamiliar mice

We next conducted a longitudinal experiment to evaluate various additional features of social threat conditioning (Fig 3A; Supp Fig 3). First, this experiment included various groups of males and females that underwent testing against male or female social stimuli in either sex-matched or sex-unmatched fashions (FvsF, N = 8; FvsM, N = 6; MvsM, N = 7; and MvsF, N = 6) to evaluate sex differences and the potential implications of opposite-sex trauma. Second, this experiment included multiple post-conditioning sociability tests at various time points against distinct unfamiliar conspecifics to evaluate the persistence of social phobia. Third, an additional sociability test was also conducted against familiar cagemates to further evaluate stimulus generalization. And fourth, multiple open field tests were intermingled within the experiment to evaluate possible generalization to non-social behaviors.

**Fig 3.**
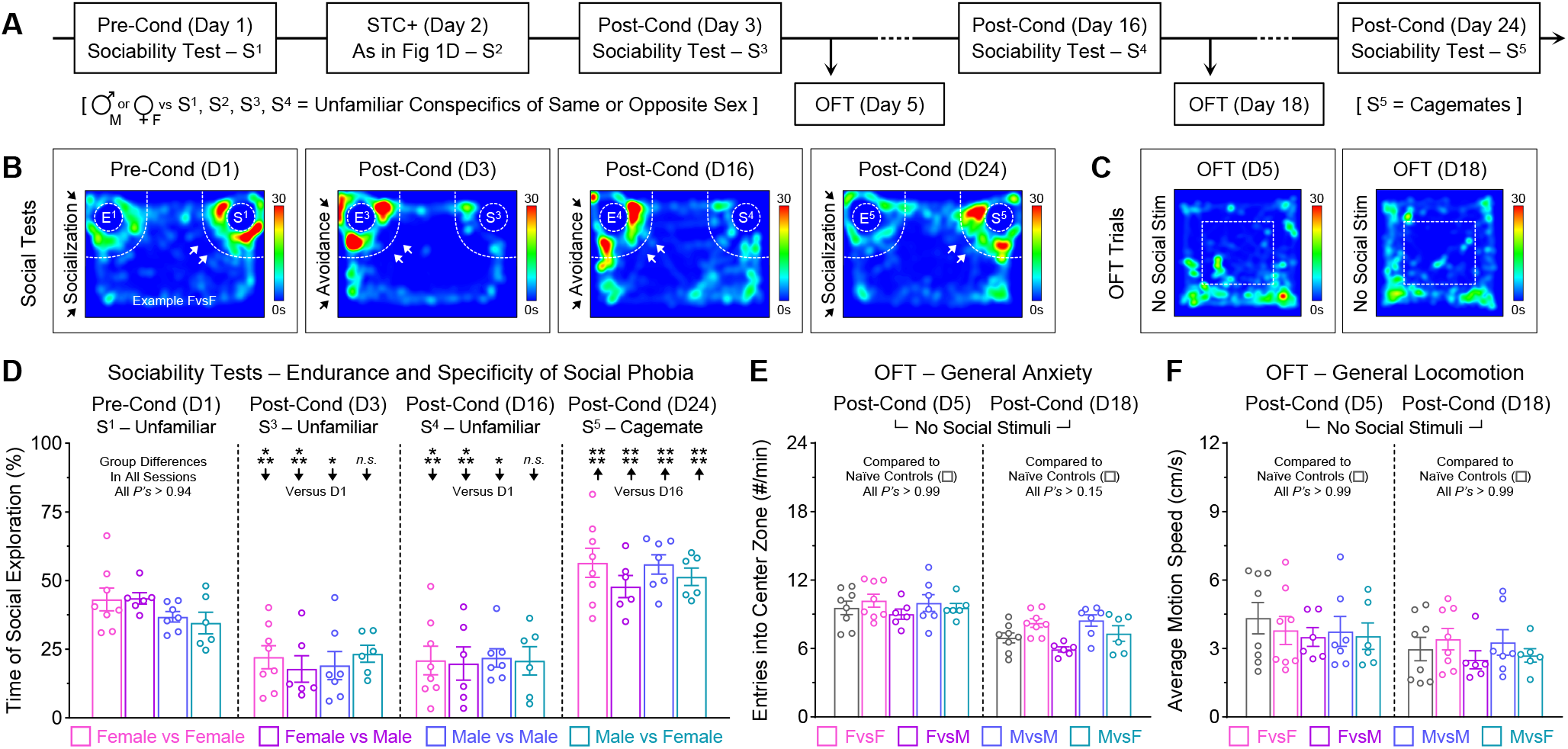
Social phobia after social threat conditioning was enduring, independent of sex variables, and particularly triggered by unfamiliar conspecifics. ***A***, Experimental design for longitudinal assessment. Sex differences and the potential influence of opposite-sex trauma were evaluated while comparing multiple groups in which the test subjects and the social stimuli had either matched or mismatched sexes (FvsF, N = 8; FvsM, N = 6; MvsM, N = 7; MvsF, N = 6; the first letter indicates the sex of the test subjects while the second letter indicates the sex of the social stimuli). ***B***, Heatmaps for a representative female subject that underwent training and testing against other females. ***C***, Heatmaps for the same representative female during open field tests. ***D***, Quantifications of social exploration during the sociability tests. One day after conditioning, all groups exhibited lower social exploration, compared to baseline (FvsF, *P* = 0.0006; FvsM, *P* = 0.0003; MvsM, *P* = 0.011; MvsF, *P* = 0.37). Fourteen days after conditioning, all groups still exhibited persistently lower social exploration, compared to baseline (FvsF, *P* = 0.0003; FvsM, *P* = 0.0008; MvsM, *P* = 0.047; MvsF, *P* = 0.13). A return in social exploration was only observed when all groups were tested against familiar cagemates a few days after (all *P’s* < 0.0001). ***E-F***, Anxiety-related behavior and general locomotion during the open field tests. Comparisons made against a naïve group (N = 8, both sexes combined). [Two-way ANOVAs with Bonferroni post-hoc tests: \**P* < 0.05, \*\**P* < 0.01, \*\*\**P* < 0.001, \*\*\*\**P* < 0.0001]

During the pre-conditioning test, regardless of sex variables, no group differences were detected in baseline sociability (Fig 3D, D1). During a post-conditioning test one day after training, while no differences were detected among the groups (Fig 3D, D3), most groups exhibited significantly lower sociability compared to the baseline session (Fig 3D, D3). Similarly, during a subsequent post-conditioning test two weeks later, while no differences were detected among the groups (Fig 3D, D16), most groups still exhibited low sociability with the unfamiliar conspecifics (Fig 3D, D16). In contrast, when tested against familiar cagemates a few days later, all groups exhibited significant rebounds in sociability (Fig 3D, D24). Finally, during the open field tests, no group differences were detected in measurements for general anxiety and locomotion, not even when compared to a naïve group (Fig 3E-F). These findings indicate that STC results in persistent social deficits, regardless of sex variables, but do not generalize to familiar cagemates or to novel environments.

### Silencing the prelimbic prefrontal cortex prevented the development of lasting social phobia

Our next goal was to test whether neural activity in the PL prefrontal region is necessary for the acquisition of memory representations of social threat. This was tested using an optogenetic silencing approach (Supp Fig 4A,C,E) in which PL was bilaterally transduced with either an inert fluorophore or a light-sensitive outward proton pump (eYFP, N = 8 males; ArchT, N = 7 males). During experiments, ArchT was triggered using red-shifted laser light, delivered through chronically implanted optical fibers.

**Fig 4.**
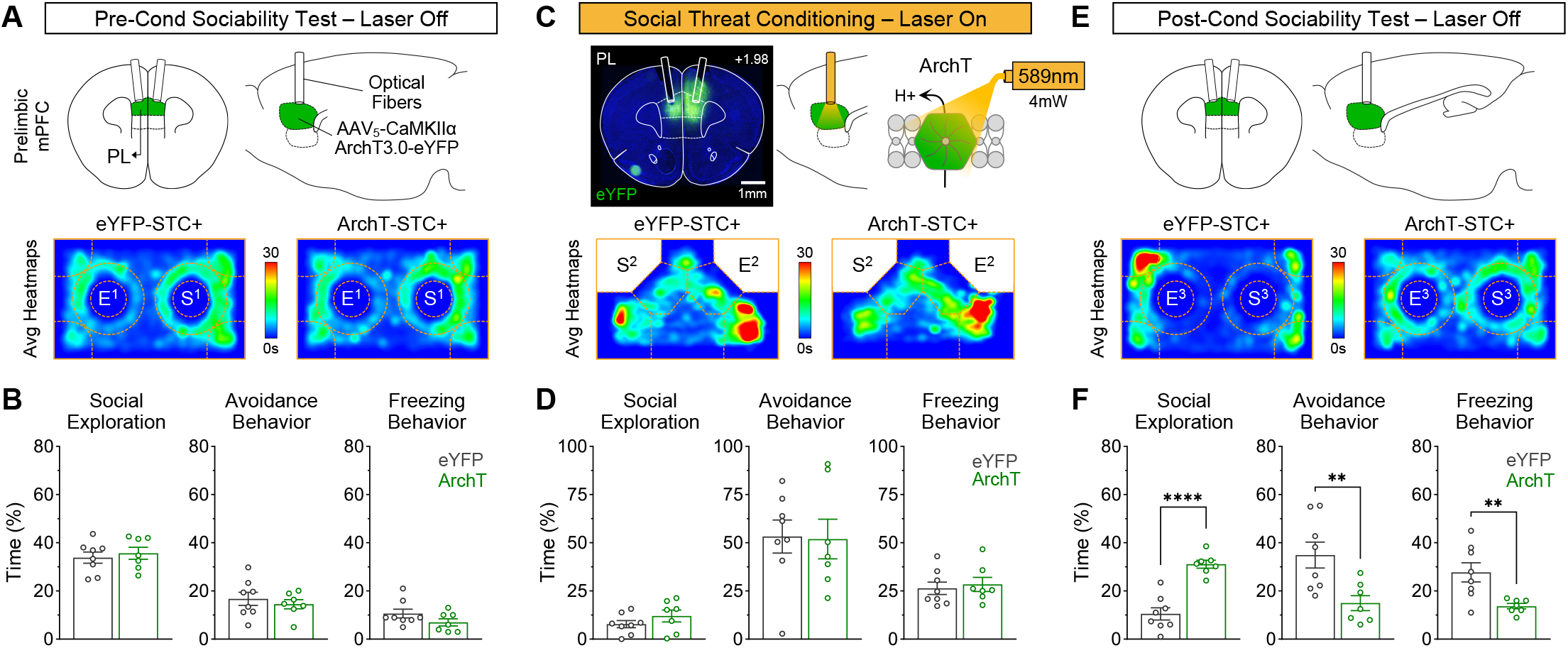
Photoinhibition of the prelimbic prefrontal cortex during social threat conditioning produced forgetting and prevented subsequent social phobia. ***A-B***, Pre-conditioning sociability test to gather the behavioral baselines for mice expressing either a control fluorophore (eYFP, N = 8 males) or an inhibitory opsin (ArchT, N = 7 males) in PL. No differences were detected between the groups during the baseline test (Social Cup, *t*_(13)_ = 0.53, *P* = 0.61; Avoidance, *t*_(13)_ = 0.62, *P* = 0.52; Freezing, *t*_(13)_ = 1.49, *P* = 0.16). ***C-D***, Social threat conditioning with optogenetic silencing of PL. Both groups exhibited reduced social exploration and increase avoidance and freezing, but no differences were detected between the groups (Social Cup, *t*_(13)_ = 1.21, *P* = 0.25; Avoidance, *t*_(13)_ = 0.10, *P* = 0.92; Freezing, *t*_(13)_ = 0.42, *P* = 0.68). ***E-F***, Post-conditioning sociability test to evaluate long-term effects. While the eYFP group continued exhibiting social phobia traits, the ArchT group exhibited high levels of social exploration and reduced avoidance and freezing, as if they did not remember the previous traumatic experience. This resulted in highly significant differences between the groups during the long-term test (Social Cup, *t*_(13)_ = 6.61, *P* < 0.0001; Avoidance, *t*_(13)_ = 3.10, *P* = 0.009; Freezing, *t*_(13)_ = 3.19, *P* = 0.007). Therefore, neural activity in PL is critical for the development of lasting social phobia. [\*\**P* < 0.01, \*\*\*\**P* < 0.0001]

As in other experiments above, the eYFP and ArchT groups underwent a three-day protocol. On day one, a pre-conditioning test was conducted without laser manipulation (Fig 4A). No group differences were detected in any of the baseline behaviors (Fig 4B). On day two, STC was conducted with laser manipulation (Fig 4C). Both groups exhibited comparable acquisition of STC (Fig 4D). On day three, a post-conditioning test was conducted without laser manipulation (Fig 4E). Despite social deficits during training, the ArchT group exhibited a behavioral profile that was strikingly similar to the behavioral profile exhibited during the initial baseline test (i.e., high social exploration and low avoidance and freezing). This resulted in significant differences when compared to the eYFP controls, which continued exhibiting robust traits of social phobia (Fig 4F). Additional experiments showed a trivial role of the adjacent IL prefrontal region (Supp Fig 4B,D,F; Supp Fig 5). Furthermore, the optogenetic treatments produced very limited effects in on-going defensive behavior during STC (Supp Fig 6) or during traditional open field testing (Supp Fig 7), which ruled out several alternative explanations. Therefore, PL (but not IL) seems critical for the acquisition of neural representations of social threat.

**Fig 5.**
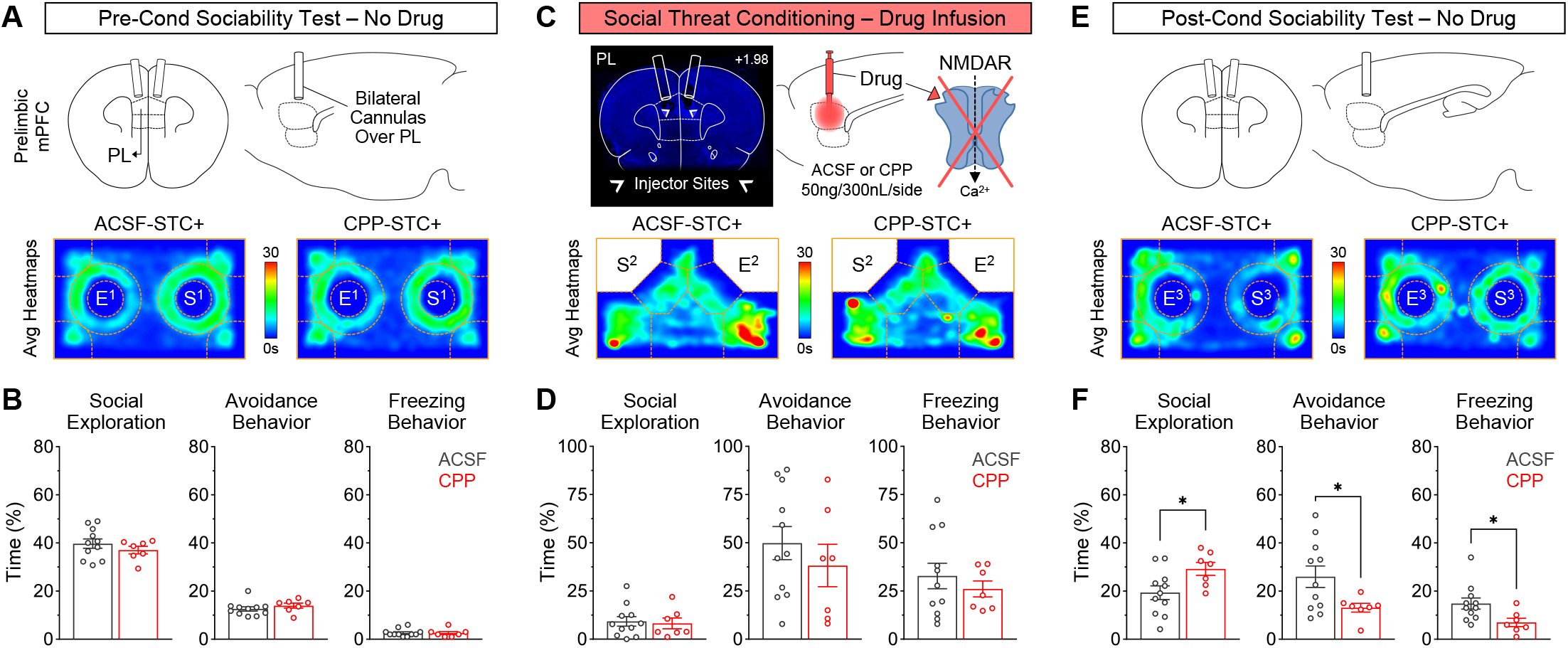
Pharmacological inhibition of NMDARs in PL also impaired lasting social phobia. ***A-B***, Pre-conditioning test for behavioral baselines. No significant differences were detected between the groups (Social Cup, *t*_(16)_ = 0.98, *P* = 0.34; Avoidance, *t*_(16)_ = 0.98, *P* = 0.34; Freezing, *t*_(16)_ = 0.08, *P* = 0.94). ***C-D***, 30 min prior to social threat conditioning, mice received bilateral infusions in PL of either vehicle (ACSF, N = 11 males) or a long-lasting competitive antagonist of NMDARs (CPP, N = 7 males). The two groups exhibited comparable social phobia traits this training session (Social Cup, *t*_(16)_ = 0.25, *P* = 0.80; Avoidance, *t*_(16)_ = 0.84, *P* = 0.41; Freezing, *t*_(16)_ = 0.74, *P* = 0.47). ***E-F***, Drug-free post-conditioning sociability test. Compared to the ACSF controls, the CPP group exhibited a significant rebound in social exploration, and significantly reduced avoidance and freezing (Social Cup, *t*_(16)_ = 2.38, *P* = 0.030; Avoidance, *t*_(16)_ = 2.22, *P* = 0.041; Freezing, *t*_(16)_ = 2.41, *P* = 0.028). These findings suggest that NMDAR-mediated plasticity in PL promotes the formation of a lasting representation of social threat. [\**P* < 0.05]

### Local inhibition of NMDARs in the prelimbic cortex also prevented lasting social phobia

While the findings above suggest that PL processing is critical for social threat learning, it remained elusive whether events related to plasticity were taking place in PL during social threat learning. To test this, a drug microinfusion approach was used to inhibit NMDARs (Supp Fig 8), which typically promote burst firing and calcium-dependent molecular cascades to facilitate synaptic plasticity and memory formation [34,35]. Thus, after placing cannulas over PL, microinfusions of either ACSF vehicle (N = 11 males) or the NMDAR antagonist CPP (N = 7 males) were performed 30 min prior to STC.

During the pre-conditioning test (Fig 5A), no differences were detected between the ACSF and CPP groups (Fig 5B). The next day, during the STC task with the drugs onboard (Fig 5C), both groups exhibited similar acquisition of social phobia traits (i.e., low social exploration, high avoidance, and high freezing), thus resulting in no group differences (Fig 5D). During a drug-free post-conditioning test the next day (Fig 5E), the CPP group exhibited significantly higher social exploration and lower avoidance and freezing than the ACSF group (Fig 5F). Therefore, NMDAR-dependent mechanisms seem to be occurring in PL to mediate the acquisition of lasting representations of social threat.

## DISCUSSION

Social threat conditioning was used in this study to provide new insights on the mechanisms by which social-related trauma leads to the development of social phobia traits in mice. At the behavioral level, our experiments revealed that this phenomenon does not depend on sex variables or the particular identity of the social predictor of threat. At the neurobiological level, our experiments showed that neural processing and NMDAR activity in the PL (but not IL) subregion of the mPFC are critical mechanisms by which social trauma leads to lasting traits of social phobia. Collectively, these findings suggest that socially-derived traumatic experiences engage learning mechanisms in the PL region to make lasting representations of social threat. These new insights could help to pave the way for future studies that evaluate potential new treatments or novel strategies for the prevention of social phobia.

### Social threat conditioning is an ideal model for studying social phobia

Despite remaining understudied, the STC task in rodents has already produced multiple striking findings, suggesting that this paradigm is ideal for studying the causes of social phobia. For instance, it has been shown that STC produces behavioral patterns of social phobia that could endure for prolonged periods without producing phobia to unanimated objects or non-social settings [15,16,36,22]. Notably, social phobia after STC is not only exhibited to the particular social stimuli that predicted shock punishment during training, but also to other unfamiliar conspecifics that had nothing to do with the previous traumatic experience, in a context independent manner [15,22]. In the present study, we replicated many of these observations, and expanded the view of social phobia after STC by showing that this phenomenon does not depend on sex variables, not even if the sex of the test subjects and the social stimuli are matched or mismatched. Furthermore, when we attempted a new version of the task that optimizes differential learning, the test subjects eventually ended up developing profiles of generalized phobia across social stimuli. The only instance in which we observed a return of social behavior back to normal levels was when the subjects were subsequently tested against familiar cagemates. Notably, these observations are consistent with the overall phenomenology observed in human patients, who mostly exhibit social phobia when they undergo interactions with unfamiliar strangers but not with close relatives [1,2]. Altogether, these observations recapitulate the notion that the STC task is ideal for studying the impact of social trauma to produce social phobia-like states, even when the task does not involve physical harm, social defeat, or aggression [11,13,37,38].

### A variety of defensive behaviors emerge during social threat conditioning

While the impacts of STC paradigm have been mostly emphasized within the realm of social behavior, in this study we described additional impacts that extend into the core of defensive behavior. For instance, animals that underwent the STC task exhibited prominent increases in anxiety-like avoidance behavior, which was not just characterized by lack of exploration of the social predictor of threat, but it was mostly characterized by full retreat into the farthest possible corner of the conditioning chamber while keeping vigilant or oriented to the social stimulus. Furthermore, the test subjects exhibited large amounts of freezing behavior, which is a robust defensive behavior in many species during threat [24,25,39]. In addition, the test subjects exhibited multiple forms of darting and stretched postures. While darting moves represents escape or flight-like responses during threat [24,25,32], stretched posture represents risk assessment and ambivalent behavior during conflict [31,40]. In this case, ambivalence most likely emerged as a consequence of conflicting drives to either avoid or approach the social stimulus. Altogether, these findings demonstrate that social threat conditioning does not only involve impacts on social behavior, but it also involves a complex repertoire of defensive behaviors that reinforce the notion of effectiveness of the task to produce strong representations of social threat. Yet, it remains to be elucidated how the individual defensive mechanisms contribute to learning during the task, and thus future studies could focus on close-loop manipulations capable of disrupting the individual behaviors.

### Social threat conditioning engages learning mechanisms within the prelimbic cortex that shape the development of lasting representations of social threat

Regarding the neurobiological bases of social threat conditioning, not much work has been done on this front. However, a recent study implicated the mPFC – particularly the PL division – in the long-term expression of social phobia after STC [22]. In that study, the expression of social phobia was associated with elevated cfos expression, increased spiking activity in principal neurons, augmented somatostatin-derived activity, and reduced parvalbumin-related activity in PL. Together, these observations suggest that disinhibitory mechanisms are engaged in PL to produce greater excitatory output to influence the expression of social phobia [13], an idea that was further tested using cell-type-specific manipulations [22]. Despite these previous observations, it remained elusive whether neuronal processing in PL during the initial social threat experience was required to shape the mechanisms that eventually produce social phobia. The present optogenetic findings are consistent with this idea, as they indicate that PL activity is indeed needed during STC training to produce lasting traits of social phobia. Furthermore, we observed that local inactivation of NMDARs in PL during STC training resulted in significant impairment in subsequent social phobia. Collectively, these observations emphasize the importance of this brain region to encode social threat during the STC task. Yet, additional studies are needed to define the neurophysiological mechanisms by which distinct populations of PL neurons encode social threat.

### What kind of information could PL neurons be encoding during social threat conditioning?

PL is an integral part of the corticolimbic networks implicated in the encoding and regulation of aversiveness and unpleasantness during threat, harm, punishment, or conflict [41–45]. Based on input connectivity from a variety of sources [46,47], the PL region is well positioned for the convergence of socially-derived and shock-related signals for the encoding of social-threat contingencies. For instance, PL receives robust glutamatergic inputs from the mediodorsal thalamus (MDT) in which large proportions of neurons get strongly activated by social cues [48,49]. Furthermore, it is thought that PL neurons are capable of decoding socially-derived signals to alter a variety of social functions, including social interaction, social preference, and social dominance [50–52]. Regarding the processing of shock-derived signals, significant proportions of PL neurons exhibit excitatory responses during shock punishment [53,54]. A main source of shock-related information to PL is believed to originate in the basolateral amygdala (BLA), which also exhibits robust excitatory signals during shock punishment and other aversive stimuli [55–60]. Nonetheless, some evidence suggests that the MDT→PL pathway could also transmit shock-related signals to PL [61], and that the BLA→PL pathway could also transmit socially-derived information to PL [62], suggesting that these pathways could already relay multisensorial signals to PL. In either case, once PL integrates these signals to represent social threat, PL could then exert top-down regulation of defensive mechanisms and social behavior to support the social phobia states [63–65]. In support of this idea, it has been shown that activation of PL outputs to the nucleus accumbens (NAc) decreases social preference [66]. Furthermore, activation of PL outputs back to BLA impairs social behavior and augments defensive mechanisms such as avoidance [67,68]. Therefore, PL appears to be well positioned not only for the encoding of social-threat contingencies, but also to support the subsequent alterations in behavior through top-down modulation of subcortical regions.

## ACKNOWLEDGEMENTS

The authors thank Stephanie Villalon, Shatavia McBride, Kamryn Whitehorn, Hope Msengi, and Carolina Gonzalez for technical assistance. The authors also thank Dr. Maria Diehl (KSU) for comments on the manuscript. A.B.R., K.L.O., A.C.F.O., J.M.T., and A.R.R. were supported by a University of Texas Program on Science and Technology Acquisition and Retention (STARs), UTSA College of Sciences, Baptist Health Foundation Award, and NIH grants R25EB027605 and T34GM145507. J.R.R. was supported by the NC TraCS Institute, the Foundation of Hope, the Brain and Behavior Research Foundation, the Kavli Foundation, the Whitehall Foundation, and the NIH grant R01MH132073.

## AUTHOR CONTRIBUTIONS

Conceptualization: A.B.R. and J.R.R. Research Design: A.B.R. and J.R.R. Methodology: A.B.R., K.L.O., A.C.F.O. Behavioral Experiments: A.B.R., K.L.O., A.C.F.O., A.R.R., J.M.T. Optogenetic Experiments: K.L.O., A.C.F.O., A.R.R. Pharmacology Experiment: A.B.R., K.L.O., A.R.R. Data Analysis: A.B.R., K.L.O., A.C.F.O. Histology: K.L.O., A.R.R., J.M.T. Supervision of all Aspects of the Study: A.B.R. Writing of Article: A.B.R., K.L.O., J.R.R.

## DATA AVAILABILITY

Data supporting this research study will be shared for appropriate scientific use upon request. Viral vector sequences and other related information could be requested from the commercial suppliers (Addgene and UNC Vector Core).

## COMPETING INTERREST

All authors declare no conflicts of interests.

