## Supplementary Materials for "The prelimbic prefrontal cortex mediates the development of lasting social phobia as a consequence of social threat conditioning"

#### SUPPLEMENTARY METHODS

##### Behavioral Analysis

ANY-maze software was used for automated quantification of most behaviors, based on mouse body center point. Social exploration was defined as the time that the test subjects spent in proximity to the cups or enclosures that contained the social stimuli. Non-social exploration was defined as the time they spent in proximity to the empty cups or enclosures. Anxiety-like avoidance was defined as the time they spent in the corners of the boxes or chambers [1,2]. Fear-related freezing was quantified as the time mice spent in complete immobility [3,4], regardless of location within the boxes or chambers. ANY-maze was also used for other quantifications, such as average entries into distinct zones, distance to social stimuli, total distance traveled, and speed of locomotion. In addition to software-based quantifications, hand-scoring methods were implemented by experimenters that remained blind to mouse treatment to evaluate additional defensive behaviors, such as darting and stretched posture [5–8]. All data was organized in spreadsheets (MS-Excel), and then analyzed and plotted using GraphPad software (Prism-10). Normality of the data was verified with the Kolmogorov-Smirnov test. The data was then plotted as mean ± sem with superimposed values for individual subjects. Unpaired T-tests were applied for analyses that only considered two groups and one dependent variable, whereas two-way repeated measures analysis of variance with Bonferroni post-hoc tests were implemented for analyses considering two dependent variables (e.g., group × time). Multiplicity adjusted $P$-values were considered if appropriate to account for multiple comparisons. The smallest statistical significance level considered was $P < 0.05$.

**Surgical Procedures**

Surgical procedures were performed under aseptic conditions to infuse viral vectors and to chronically implant optical fibers or cannula probes. Surgeries were performed using stereotaxic frames (KOPF) that were equipped with isoflurane gas vaporizers for anesthesia (Harvard Apparatus). Pre-emptive analgesia was provided through local application of lidocaine and bupivacaine (7-8 mg/kg each) and systemic injections of sustained-release meloxicam (Melox-SR, 4 mg/kg, s.c.). Skin incisions were then opened, and craniotomies were drilled using precision microdrills (35k RPM; Alkita) and tungsten carbide drill bits (0.5-0.7 mm tips; Fisher Scientific). PL and IL were targeted bilaterally and individually as in a previous study [9]. Stereotaxic coordinates for viral infusions in PL: +1.70 mm A/P, ±0.35 mm M/L, and -2.40 mm D/V. Stereotaxic coordinates for optical fibers in PL: +1.70 mm A/P, ±0.75 mm M/L, -1.95 mm D/V, at a 10° M/L angle. Stereotaxic coordinates for viral infusions in IL: +1.73 mm A/P, ±0.35 mm M/L, and -2.95 mm D/V. Stereotaxic coordinates for optical fibers in IL: +1.73 mm A/P, ±1.25 mm M/L, -2.30 mm D/V, at a 20° M/L angle. Stereotaxic coordinates for guide cannulas in PL: +1.70 mm A/P, ±0.75 mm M/L, -1.95 mm D/V, at a 10° M/L angle. All the coordinates above were relative to bregma. Optical fibers and cannulas were then secured to the skull using biocompatible adhesive cement (C&B Metabond; Parkell) and black orthodontic acrylic (Ortho-Jet; Lang Dental), and incisions were finally sutured with black nylon (4-0; Covetrus). Mice were then transferred to clean cages with regulated temperature and free access to soft food (DietGel 76A; Clear $H_2O$). After a few hours, mice were brought back to the vivarium for full recovery. Post-operative care was provided by the experimenters and veterinarian staff for at least 5 days. If necessary, additional injections of Melox-SR and lactate ringers were administered.

**Optogenetic Silencing**

Viral vectors were obtained from a commercial source (UNC Vector Core) to express either enhanced yellow fluorescent protein in the control groups ($AAV_5$-CaMKIIα-eYFP) or the light-sensitive outward proton-pump archaerhodopsin in the experimental groups ($AAV_5$-CaMKIIα-ArchT3.0-eYFP). Viral infusions were performed during surgeries using 10-µL Nanofil syringes and 33-Ga needles that were driven with an ultramicropump (UMP3T-2; WPI). A volume of ~400 nL of the virus-containing medium was infused in each hemisphere, at a rate of 100 nL/min. The infusion needles were then kept at the target sites for an additional 10 min, after which they were slowly withdrawn. This was followed by chronic implantation of optical fibers, which were constructed in-house using pure-silica hard-cladding multimode fibers (Ø300-µm core, NA = 0.39, Low OH; Thorlabs), epoxied to 304 stainless-steel ferrules (1.25 mm OD, 330 µm ID bore; Precision Fiber Products). Optical fibers were then micropolished (SpecPro; Krell Tech) until achieving light transmission >85%, measured with a light power meter (PM100D; ThorLabs). During experiments, using ceramic split sleeves (Ø1.25-mm; Precision Fiber Products), the optical fiber implants were connected to branching fiberoptic patch cords (1x2, Ø200-µm core, FCM-2xMF1.25; Doric Lenses) and fiberoptic rotary joints (FRJ-1x1-FC-FC; Doric Lenses), which were in turn connected to red-shifted DPSS lasers (589-nm wavelength; OptoEngine) using FC/PC patch cables (Ø200-µm core, NA = 0.22, Low OH; ThorLabs). Laser beam output was moderated using mechanical shutters (100-Hz SR475; SRS), which were controlled with TTL signals that were provided by an optogenetic computer interface (ANY-maze). During experiments, laser light was delivered in a constant fashion at a power of ~4-6 mW (~14-22 $mW/mm^2$), as animals underwent social threat conditioning.

**Drug Microinfusions**

For microinfusing drugs into PL, cannula systems were fabricated in-house using 304-hypodermic stainless-steel tubing (Small Parts). The fabricated parts included cannula guides that provided a clear path into the brain targets (25-Ga, 500 µm OD, 360 µm ID, cut at 5 mm from pedestal), cannula dummies that were cut to the same length as the cannula guide tubes (30-Ga, Ø280 µm), and cannula injectors that extended 500 µm from the tip of the guide tubes to reach the targets (32-Ga, 230 µm OD, 130 µm ID). In two animals from the control group, the injectors extended 1000 µm because the cannula guides were implanted at a higher position. At least 3 days prior to drug infusions, dummies were removed from the guide tubes, and clean unloaded injectors were inserted to prime the injector tracks into the tissue. During experiments, the injectors were loaded with ACSF or CPP, and were connected to 10-µL Nanofil syringes (WPI) via fluorinated ethylene-propylene tubing (680 µm OD, 120 µm ID; Sci Pro). The syringe plungers were then lowered using an ultramicropump (UMP3T-2; WPI), to infuse drugs at a rate of 100 nL/min ~30 min prior to the beginning of the social threat conditioning session.

#### Euthanasia and Histology

Mice were deeply anesthetized with isoflurane and transcardially perfused using ice-cold 1X-PBS and 4%-PFA. Brains were then collected, fixed for an additional 24-hr period in 4%-PFA, and further equilibrated in 30% sucrose. Coronal sections were then cut at 40 μm using a microtome (HM430; Thermo Scientific). Coronal sections containing the mPFC were mounted on superfrost-plus slides and cover-slipped using a fluorescence-compatible mounting medium with DAPI (Fluoromount-G; Southern Biotech). The mPFC was then imaged using a fluorescence microscope that included a software-controlled scanning stage and a high-resolution DP74 20.7MP camera for automated image stitching (BX43, Cell Sens; Olympus). The shape and transitions of cortical layers, overall tissue landmarks, and other cytoarchitectonic features were visually inspected to determine the anatomical boundaries between the PL and IL regions [10,11]. Viral expression, optical fiber placements, and cannula positions were finally evaluated and reconstructed onto coronal drawings adapted from a mouse brain atlas [12]. Several subjects were excluded from the study (21/75, 28%) due to lack of viral expression, viral leakage, or mistargeting of the PL and IL regions with either the optical fibers or cannula probes.

### SUPPLEMENTARY RESULTS

#### New hardware to perform social threat conditioning in a fully automated manner.

Social threat conditioning was initially developed in a rudimentary manner in which social contacts between test subjects and social stimuli were visually inspected by a human experimenter who eventually triggered the footshocks manually to the test subjects [13,14]. While this strategy could introduce significant variability and inconsistencies, the first goal of this study was to develop new hardware that allows to perform social threat conditioning in fully automated manner using computer-controlled operant conditioning chambers. For this, we designed novel cage enclosures to confine the animals that served as social stimuli for their presentation to the test subjects within the conditioning chambers.

The cage enclosures were designed using 3D-CAD freeware, and then fitted with acrylic panels on the ceiling and stainless-steel wire-mesh screen on the front opening to provide a way through which the test subject and the social stimulus could interact with each other (Supp Fig 1A). The enclosures were also equipped with infrared photodetectors on both sides of the mesh screen for automated detection of physical interactions between the mice (e.g., nose-to-nose contacts). Simultaneous activation of the photodetectors on both sides of the screen for at least 1 s resulted in footshocks to the test subjects. Two of these enclosures were added to congruent corners in each chamber. While one enclosure was used for presenting the social stimulus that predicted footshock, the other enclosure was kept empty in most experiments to provide additional measurements (e.g., non-social exploration).

The effectiveness of these cage enclosures was tested in the initial experiment, in which during the social threat conditioning phase, the control group did not receive footshocks during social interactions (STC-, N = 10), whereas the experimental group received footshocks during social interactions (STC+, N = 10). After 20 min, the STC+ group exhibited a significantly lower number of social bouts with the unfamiliar conspecific, compared to the STC- group (Supp Fig 1B, Left Panel, $t_{(18)} = 3.20$, $P = 0.005$). Furthermore, the STC+ group remained at a significantly higher distance to the unfamiliar conspecific, compared to the STC- group (Supp Fig 1B, Right Panel, $t_{(18)} = 3.92$, $P = 0.001$). These findings are consistent with significant reductions in social behavior and significant avoidance of the social stimulus. Therefore, the novel cage enclosures are highly effective to achieve social threat conditioning.

**Additional defensive mechanisms observed during social threat conditioning.**

In the main article, it was demonstrated that social threat conditioning produced robust reductions in social exploration, as well as significant increases in avoidance and freezing behaviors (Fig 1), which represent passive mechanisms for safety and survival [3,4]. Yet, while visually inspecting representative videos, it was evident that other defensive mechanisms also emerged during conditioning (Supp Fig 2). For instance, during social avoidance, the STC+ group exhibited a significant increase in stretched postures, which represent risk assessment and avoidance-approach ambivalence during threat [7,15]. In addition, after receiving the first footshock, the STC+ group exhibited a significant increase in darting moves, which represent active escape and flight-like responses that are critical for safety and survival [6,16]. Interestingly, further inspection revealed that relative to the direction of movement, multiple subtypes of darting were identified, including forward darting moves, reverse darting moves, and stimulus-oriented darting moves, all of which were most often performed to gain more distance from the social stimulus or predictor of threat. The emergence of these defensive behaviors further reinforces the notion that social threat conditioning produces robust representations of social threat.

**During the longitudinal experiment, no group differences were apparent in any of the sessions.**

While heatmaps for a representative test subject were provided in the main article (Fig 3B-C), for full transparency of the results all the group heatmaps per session are provided in these supplementary materials (Supp Fig 3). As an additional important note, the sociability tests during this experiment included a slight modification to the positioning of the inverted cups, which were located closer to two congruent corners of the apparatus. While no clear benefits were gained with this modification, it was clearly evident that after social threat conditioning all groups tended to use the space between the empty cup and the walls of the apparatus to avoid the social stimulus (Supp Fig 3, D3 and D16). Despite this, a similar outcome was obtained in all groups (i.e., deficits in social behavior), regardless of sex variables.

**Strategy and histology for the experiments involving optogenetic silencing of prefrontal regions.**

Principal neurons in either the prelimbic (PL) or infralimbic (IL) subregions of the mPFC were transduced with either a fluorophore (eYFP) or a light-sensitive outward proton pump (ArchT). The opsin was activated with laser light, delivered through chronically implanted optical fibers. Full reconstructions for the position of optical fibers and the centroid of viral expression within the PL or IL are provided in the supplementary figures below (Supp Fig 4).

**Silencing the infralimbic cortex did not prevent the development of lasting social phobia.**

In the main article, it was shown that neural silencing in the PL subregion of the mPFC produced significant forgetting and abolishment of lasting social phobia (Fig 4). In addition to PL, the IL subregion of the mPFC also plays critical roles in regulating threat-related behavior, which has been mostly demonstrated using non-social threat conditioning tasks [9,17–22]. Thus, an additional optogenetic silencing experiment was conducted to evaluate any possible role of the IL region during social threat conditioning. For this, two group of mice were prepared, which bilaterally expressed either the inert control fluorophore only (eYFP, N = 11 males) or the neuronal silencing opsin (ArchT, N = 10 males).

The role of IL was evaluated using a three-day protocol, similar to the PL experiment. During the pre-conditioning test (Fig 5A), no significant differences were detected between the eYFP and ArchT groups in the baseline level of either of the behaviors (Fig 5B; Social Cup, $t_{(19)} = 1.38$, $P = 0.18$; Avoidance, $t_{(19)} = 0.88$, $P = 0.39$; Freezing, $t_{(19)} = 1.31$, $P = 0.20$). During social threat conditioning (Fig 5C), both groups exhibited similarly robust acquisition of social phobia (i.e., exhibited low levels of social exploration, high levels of avoidance-related behavior, and high levels of freezing behavior), and no significant differences between the groups were shown by the statistical tests (Fig 5D; Social Cup, $t_{(19)} = 0.91$, $P = 0.38$; Avoidance, $t_{(19)} = 1.15$, $P = 0.26$; Freezing, $t_{(19)} = 0.21$, $P = 0.84$). During the post-conditioning test (Fig 5E), both the eYFP and the ArchT continued to exhibit traits of social phobia, and no significant differences were detected between the groups (Fig 5F; Social Cup, $t_{(19)} < 0.01$, $P > 0.99$; Avoidance, $t_{(19)} = 1.60$, $P = 0.13$; Freezing, $t_{(19)} < 0.01$, $P > 0.99$). These results suggest that the IL prefrontal region does not play a significant role for the acquisition of lasting representations of social threat.

**Silencing of the prefrontal regions produced limited effects in on-going defensive behavior.**

It was noted in the first experiment that in addition to alterations in social, avoidance, and freezing behaviors, social threat conditioning also elicited defensive mechanisms such as darting and stretched postures. Prefrontal manipulations could alter these behaviors [21,23–25]. We thus evaluated whether our optogenetic treatments produced alterations in on-going defensive behavior (Supp Fig 6A) that could possibly explain the subsequent impairments in lasting social phobia.

Optogenetic silencing of either PL or IL did not produce significant effects in any of the darting moves (Supp Fig 6B; eYFP vs ArchT; PL Darting, all *P's* > 0.48; IL Darting, all *P's* > 0.22). In contrast, stretched postures seemed particularly sensitive to PL manipulations, but not IL manipulations. Particularly, the PL-ArchT group exhibited a significant elevation in cumulative time for body stretching compared to the PL-eYFP group (Supp Fig 6B; PL Body Stretching, $t_{(13)} = 2.41$, $P = 0.032$; IL Body Stretching, $t_{(19)} = 1.00$, $P = 0.33$). This is an intriguing finding because the PL silencing treatment did not produce alterations in the other behaviors that were mainly investigated during the social threat conditioning phase. Since the PL treatment eventually resulted in forgetting and prevention of lasting social phobia, it remains to be determined whether the increases in stretch postures have something to do with the processing of social threat or whether the increases in stretch postures just reflect trivial secondary effects.

**The optogenetic treatments produced no significant effects during traditional open field testing.**

Additional experiments were conducted in the optogenetic groups above. In particular, we wanted to rule out the possibility of secondary effects, such as general anxiety and locomotion, which have been linked to mPFC activity [26,27]. To explore these possibilities, we implemented traditional open field testing (i.e., without presenting social stimuli), and included Laser-Off and Laser-On epochs (Supp Fig 7A). As proxy for general anxiety, quantifications were made for total time spent in the center of the arena [28,29]. In the PL groups, no significant effects were detected on time in the center (Supp Fig 7B; Group, $F_{(1,13)} = 0.69$, $P = 0.42$; Laser Epoch, $F_{(2,26)} = 2.05$, $P = 0.17$; Interaction, $F_{(2,26)} = 0.96$, $P = 0.40$). In the IL groups, some significant effects were detected on time in the center, but post-hoc tests revealed no significant differences between the group during any particular laser epoch (Supp Fig 7C; Group, $F_{(1,19)} = 5.70$, $P = 0.28$; Laser Epoch, $F_{(2,38)} = 2.23$, $P = 0.15$; Interaction, $F_{(2,38)} = 0.82$, $P = 0.45$; Bonferroni post-hoc tests, all *P's* > 0.078). As proxy for general locomotion, quantification were made for total distance traveled within the arena [28,29]. In the PL groups, some trends were exhibited on total distance traveled, but post-hoc tests revealed no significant differences between the group during any particular laser epoch (Supp Fig 7D; Group, $F_{(1,13)} = 0.04$, $P = 0.84$; Laser Epoch, $F_{(2,26)} = 3.13$, $P = 0.065$; Interaction, $F_{(2,26)} = 2.78$, $P = 0.081$; Bonferroni post-hoc tests, all *P's* > 0.24). In the IL groups, no significant effects were detected on total distance traveled (Supp Fig 7E; Group, $F_{(1,19)} = 0.34$, $P = 0.57$; Laser Epoch, $F_{(2,38)} = 2.94$, $P = 0.074$; Interaction, $F_{(2,38)} < 0.01$, $P > 0.99$). These results rule out possible alternative explanations for the main effects, such as modulation of general anxiety and/or locomotor activity.

**Strategy and full histology for the experiment involving drug microinfusions.**

Guide cannulas were implanted over PL, through which ACSF or the NMDAR antagonist CPP were infused to evaluate the necessity of plasticity-related events during the encoding of social threat. While a histology sample is provided in the main article, an additional example and a full reconstruction of cannula placements and microinfusion sites is provided in the supplementary figures below (Supp Fig 8).

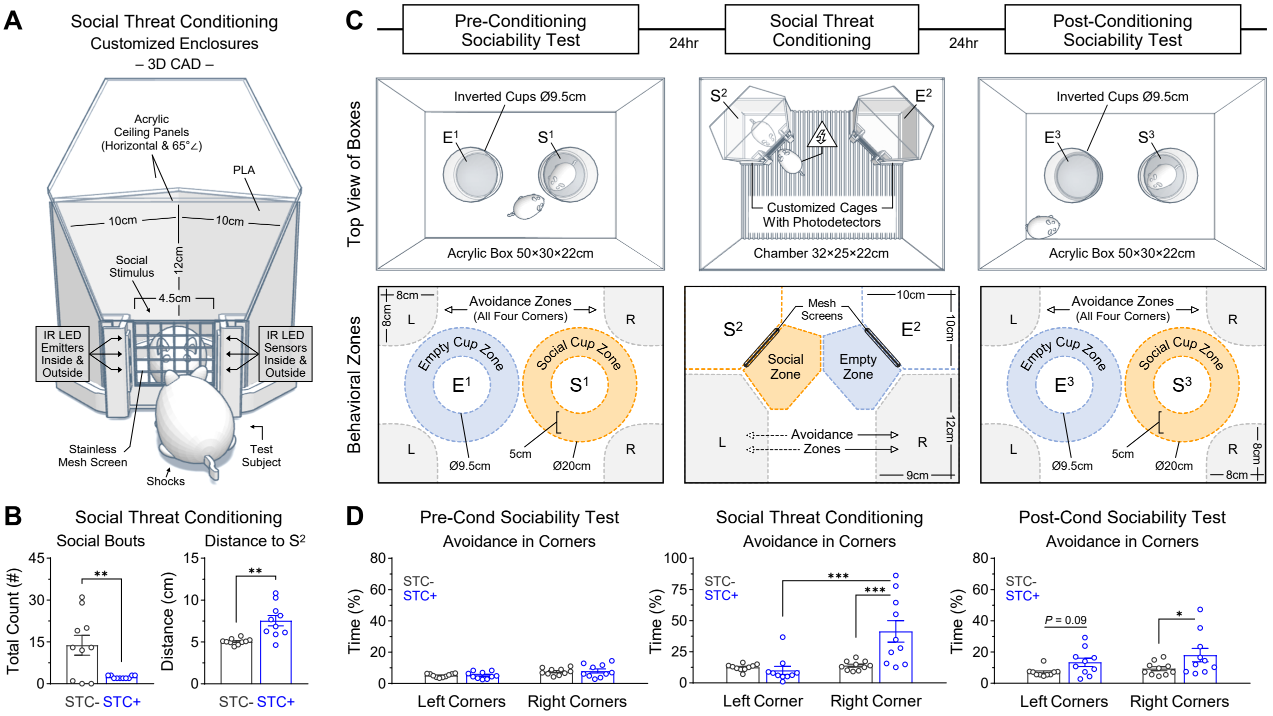


**Supp Fig 1.** Features and dimensions of hardware, depiction of the zones used for behavioral quantifications, and supplementary measurements during the initial validation of the social threat conditioning task. ***A,*** Illustration of the enclosures used during social threat conditioning. These enclosures were customized in-house and included infrared photodetectors on both sides of the mesh screen for automated detection of animals. Simultaneous activation of the photodetectors on both sides of the mesh screen indicated coincidence of social engagement, which if it lasted for at least 1 s, an electric footshock (0.5 s, 0.4 mA) was delivered to the test mouse. ***B,*** Measurements of social bouts and distance to the social stimulus are provided for animals that received shocks upon approach to the social stimulus (STC+, N = 10) and controls that did not receive shocks upon approach to the social stimulus (STC-, N = 10) during a social threat conditioning phase. ***C,*** Schematics for the full task, including the sociability tests that occurred one day before and one day after social threat conditioning. The bottom panels depict the distinct zones in each apparatus that were used for quantifications of the distinct behaviors. ***D,*** Comparisons of avoidance behavior in the left versus right corners of the boxes during each phase of the experiment. During the conditioning phase, that animals that received shocks performed more robust avoidance by retreating into the farthest possible corner of the chamber. [Unpaired T-tests in panel *B*: $^{**}P < 0.01$; two-way ANOVAs in panel *D*: $^{*}P < 0.05$, $^{***}P < 0.001$]

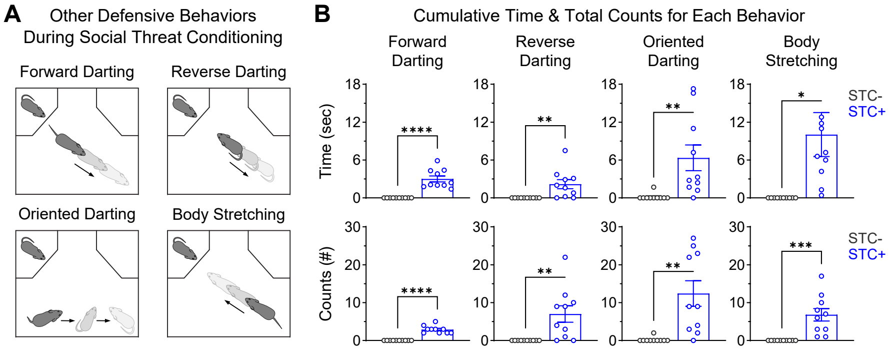


**Supp Fig 2.** Defensive behaviors related to escape and avoidance-approach ambivalence also emerged during social threat conditioning. ***A,*** Schematics for the most notable behaiors. ***B,*** Quantifications for these behaviors. [STC-, N = 10 males; STC+, N = 10 males; Unpaired T-tests: *$P < 0.05$, **$P < 0.01$, ***$P < 0.001$, ****$P < 0.0001$]

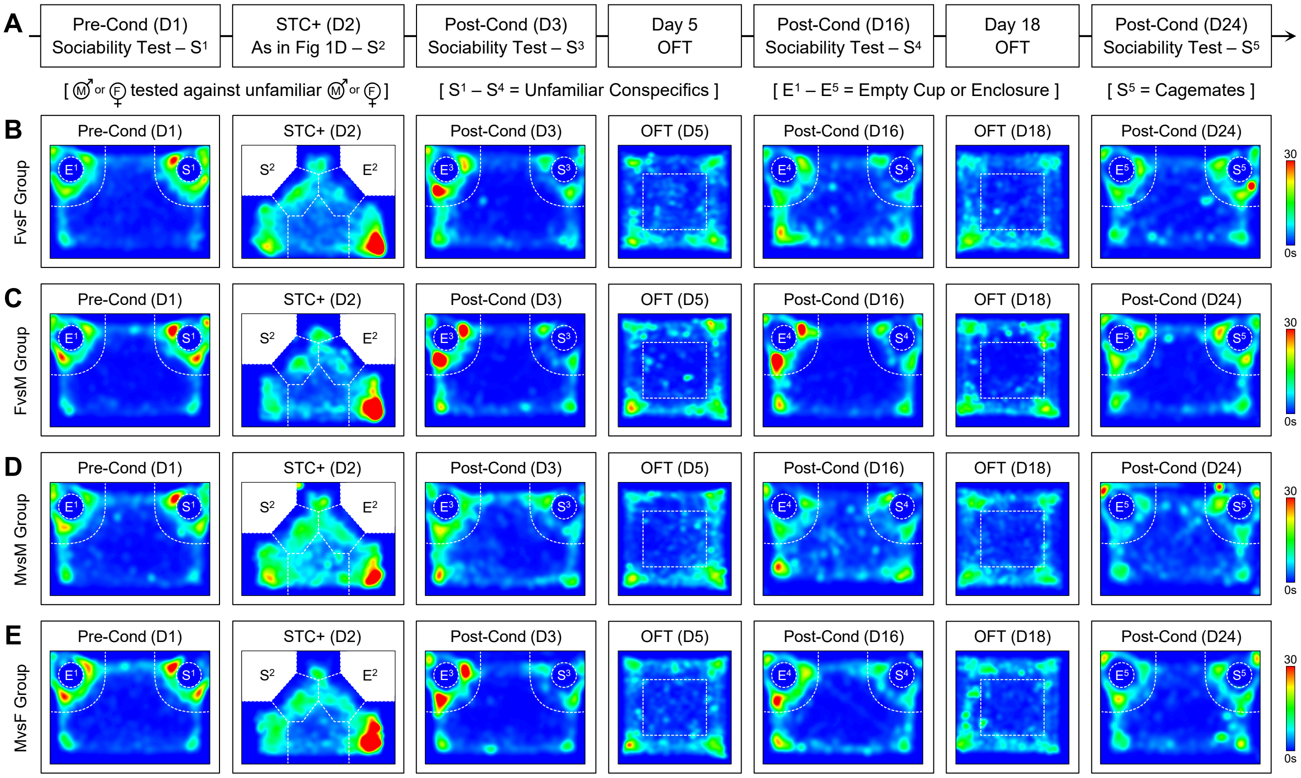


**Supp Fig 3.** Group heatmaps for all phases during the longitudinal experiment (Fig 2). ***A,*** Schematic of the experimental timeline. ***B,*** Female subjects that received shocks during exploration of unfamiliar females (FvsF, N = 8). ***C,*** Female subjects that received shocks during exploration of unfamiliar males (FvsM, N = 6). ***D,*** Male subjects that received shocks during exploration of unfamiliar males (MvsM, N = 7). ***E,*** Male subjects that received shocks during exploration of unfamiliar females (FvsF, N = 6). Mouse tracking was performed using body center as the reference point.

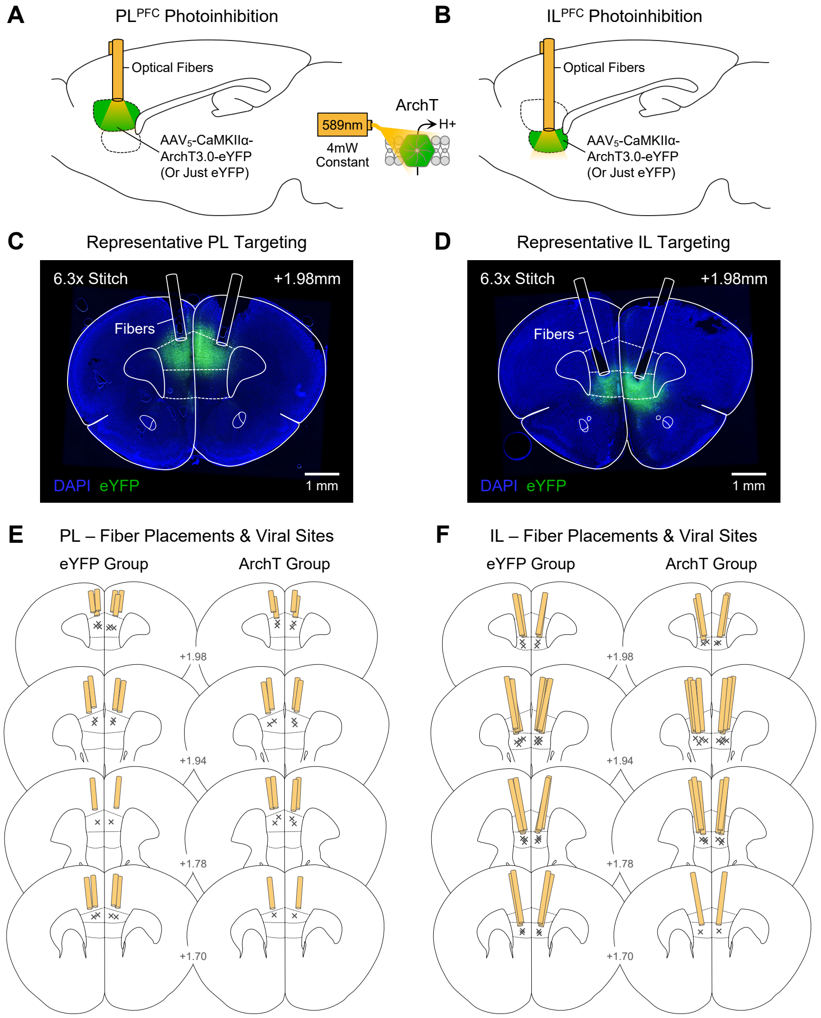


**Supp Fig 4.** Histology for optogenetic experiments. ***A-B,*** Strategy to inhibit principal neurons in either the PL or IL subregions of the mPFC. ***C-D,*** Representative coronal sections showing the targeting of PL or IL. ***E-F,*** Full reconstruction of the position of optical fibers (yellow tubes) and centroid of viral expression ("×" symbols). [PL-eYFP, N = 8 males; PL-ArchT, N = 7 males; IL-eYFP, N = 11 males; IL-ArchT, N = 10 males]

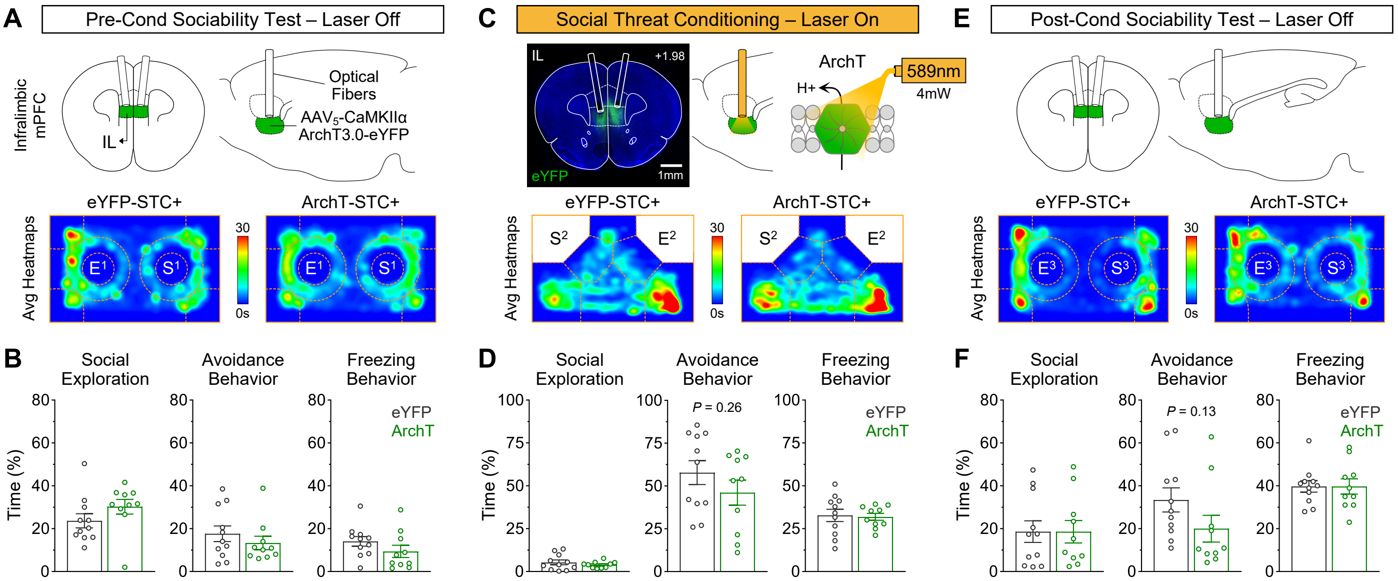


**Supp Fig 5.** Photoinhibition of the infralimbic cortex did not prevent the emergence of lasting social phobia. ***A-B,*** Pre-conditioning sociability test to gather behavioral baselines for mice expressing either a control fluorophore (eYFP, N = 11 males) or an inhibitory opsin (ArchT, N = 10 males) in CaMKII+ cells in the IL region. ***C-D,*** Social threat conditioning during optogenetic manipulation. Both groups exhibited strong behavioral traits of social phobia. ***E-F,*** Post-conditioning sociability test. Both groups continued to exhibit social phobia. This suggests that IL activity is trivial for the acquisition of prolonged representations of social threat and the emergence of lasting social phobia. [Unpaired T-tests: all *P's* > 0.05]

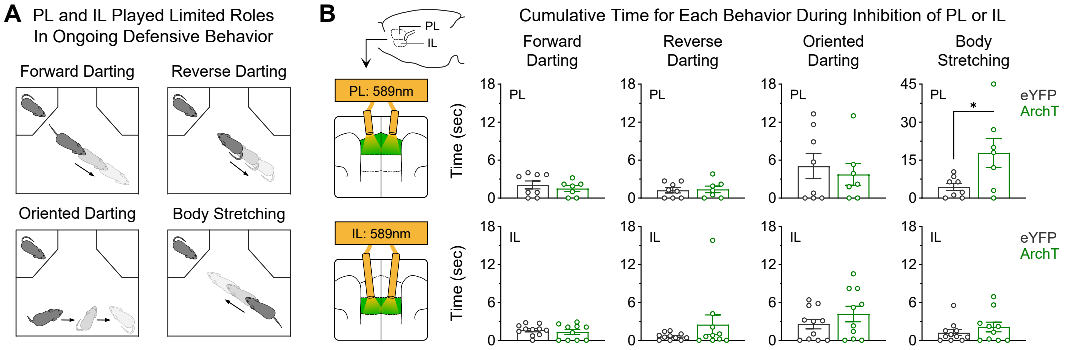


**Supp Fig 6.** The PL and IL treatments produced very limited effects in the behaviors associated with escape and avoidance-approach ambivalence. ***A,*** Schematics for the defensive behaviors of interest. ***B,*** Quantifications per behavior during photoinhibition. Only body stretching exhibited significant alterations with PL inhibition. [PL-eYFP, N = 8 males; PL-ArchT, N = 7 males; IL-eYFP, N = 11 males; IL-ArchT, N = 10 males; Unpaired T-test: *$P < 0.05$]

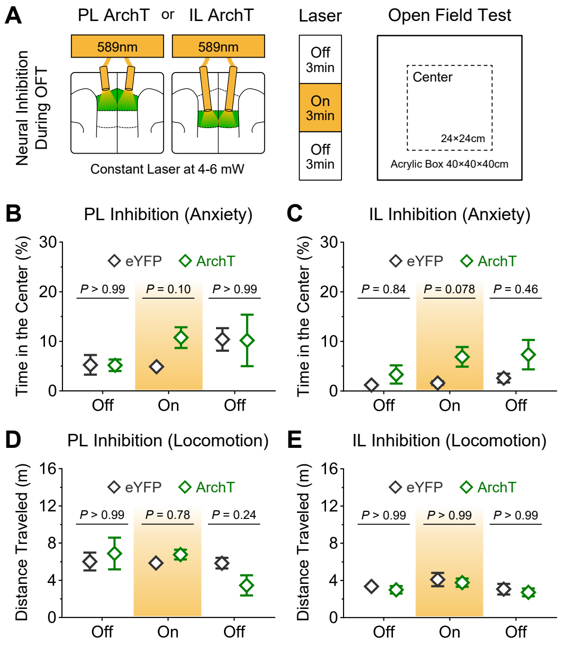


**Supp Fig 7.** PL and IL silencing produced no significant effects in general anxiety and locomotion. ***A,*** Schematics for the experimental design. Time spent in the center of the open field arena was used to assess anxiety-related behavior. ***B-C,*** The photoinhibition treatments tended to increase time in the center, but these effects did not reach statistical significance. ***D-E,*** The photoinhibition treatments did not affect locomotor activity. [PL-eYFP, N = 8 males; PL-ArchT, N = 7 males; IL-eYFP, N = 11 males; IL-ArchT, N = 10 males; *P*-values represent two-way ANOVA tests with Bonferroni post-hocs]

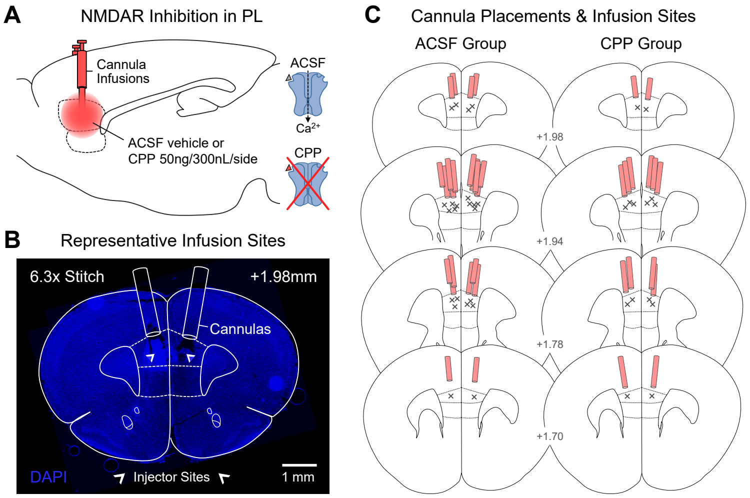


**Supp Fig 8.** Histology for the drug infusion experiment. ***A,*** Bilateral guide cannulas were chronically implanted over the PL region, and drugs were delivered through cannula injectors that extended 500 µm from the guide tubes. ***B,*** Representative cannula tracks and microinfusion sites in PL. ***C,*** Full histological reconstruction for the guide cannula tracks (red tubes) and infusion sites ("×" symbols). [ACSF, N = 11; CPP, N = 7]
